# Bioreactor-Enabled Extracellular Vesicle Production for Downstream Functional Engineering

**DOI:** 10.1101/2025.11.21.689817

**Authors:** Thaddeus L Tripp, Karina Vasile, Massimo Terrizzi, Ethan Cisneros, Lisa Volpatti, Colin Hisey

## Abstract

Extracellular vesicles (EVs) hold significant therapeutic potential, yet clinical translation is hindered by production yield and low-efficiency post-isolation engineering methods, highlighting the need for high-density bioreactor systems that support both reliable production and downstream modification. Here, we systematically compare EV output from standard flask cultures, a membrane-based two-chamber bioreactor, and a hollow-fiber bioreactor. Across culture conditions, hollow-fiber bioreactors produced markedly higher EV concentrations while requiring significantly less culture medium, demonstrating major advantages in scalability and resource efficiency. EV production was assessed with nanoparticle tracking analysis, immunoblotting, and transmission electron microscopy. Additionally, proteomic analysis revealed that bioreactor-derived EVs maintained canonical phenotypes and translational feasibility in the context of post-isolation EV engineering. Engineering feasibility was assessed with the first known reported instance of cargo-loading through EV-micelle hybridization, surface-modification through post-insertion of molecules carried by micelles (MCs), and exploitation of Nanoparticle Tracking Analysis (NTA) detection limits. Bioreactor-derived EVs remained readily engineerable, and all engineering experiments required only a small fraction of total EVs produced. Together, these findings demonstrate that bioreactor platforms overcome critical throughput limitations of conventional cultures while producing engineerable EVs. This integrated assessment establishes hollow-fiber and membrane-based bioreactors as scalable, translation-oriented systems for improved EV production and drug delivery potential via post-production engineering.

## Introduction

Extracellular vesicles (EVs) are broadly defined as lipid bilayer–enclosed particles made by cells that carry proteins, lipids, and nucleic acids, and are known to be active participants in both physiological and pathological signaling pathways (1–4). For instance, inhibition of EV-uptake via CD9 and CD63 blocking on EVs secreted by cancer cells has been demonstrated to significantly regress tumor metastasis (5); and stem cell-derived EVs have been shown to promote cell migration, proliferation, and angiogenesis during wound healing (6–9). Such intrinsically diverse biological functionality and compatibility present immense potential for regenerative medicine, targeted drug delivery, and vaccine development, and has thus garnered significant research interest (10–13).

To move EV research closer to clinical application, post-production EV-engineering approaches (broadly categorized into cargo-loading or surface modification) are leveraged to enhance specificity, efficacy, and functional payload (14,15). For example, EVs have been surface-modified via ligand conjugation to serve as targeted delivery vectors in cerebral ischemia therapies (16), and Doxorubicin has been loaded into EVs by way of electroporation to inhibit tumor growth (17,18). Nevertheless, the implementation of these clinical innovations is dependent on overcoming the foundational challenge of consistent, efficient, and scalable production of EVs, as current methods for EV engineering yield quantities insufficient for sustained clinical translation (18–20). Indeed, the current gold-standard for EV-production utilizes two-dimensional culture flasks (CF), which are severely constrained by surface area and resource intensity (14,21–24). Thus, there is a need to precede EV engineering with high-throughput EV production that is compliant with current Good Manufacturing Practices (cGMP).

Bioreactors have emerged as a superior alternative to CF due to the potential for continuous harvesting, high-density cultures, and reduced resource consumption (25–30). Critically, bioreactors have been demonstrated to produce EVs at significantly improved yields compared to CF (31,32). It is important to note, though, that a translational outlook on EV production calls for bioreactor platforms that avoid inducing stress-related phenotypes that could alter natural EV composition, while also maintaining conditions that are conducive to downstream EV engineering (33–35). Employed bioreactor platforms should then introduce minimal stress to cells and maintain the high cell densities found in vivo. Systems meeting these criteria include two-chamber diffusion bioreactors (herein referred to as CELLine bioreactors, CLB), and hollow-fiber bioreactors (HFB) due to their minimal mechanical perturbation on cells. However, these platforms have not yet been well characterized or benchmarked against each other or CF EV production. Additionally, demonstration of compatibility between bioreactor-derived EVs and post-production EV engineering workflows remains an incomplete step in establishing scalable, translatable, cGMP-compliant EV platforms. There is therefore a need to characterize CF, CLB, and HFB EV production in terms of yield, size distribution, surface markers and cargo, morphology, and engineerability.

This study utilizes CLB, HFB, (depicted in Figure 1A) and CF systems to directly compare particle size and concentration, marker expression, proteomic profiles, and morphology associated with respective EV production. Human embryonic kidney cells (HEK293T) were used to produce EVs due to their prevalence in biopharmaceutical engineering (36,37). We focus on elucidating the engineering-amenability of bioreactor-derived EV production using incubation and sonication to surface-modify and cargo-load EVs with representative, fluorescent molecules vectorized by micelles (MCs). To the best of our knowledge, we are the first to leverage the detection limit of Nanoparticle Tracking Analysis to verify EV-MC hybridization and present the first demonstration of EV-MC hybridization. This study further provides a framework for cGMP oriented, scalable EV production that supports both functional engineering and downstream translational applications.

**Figure 1.**
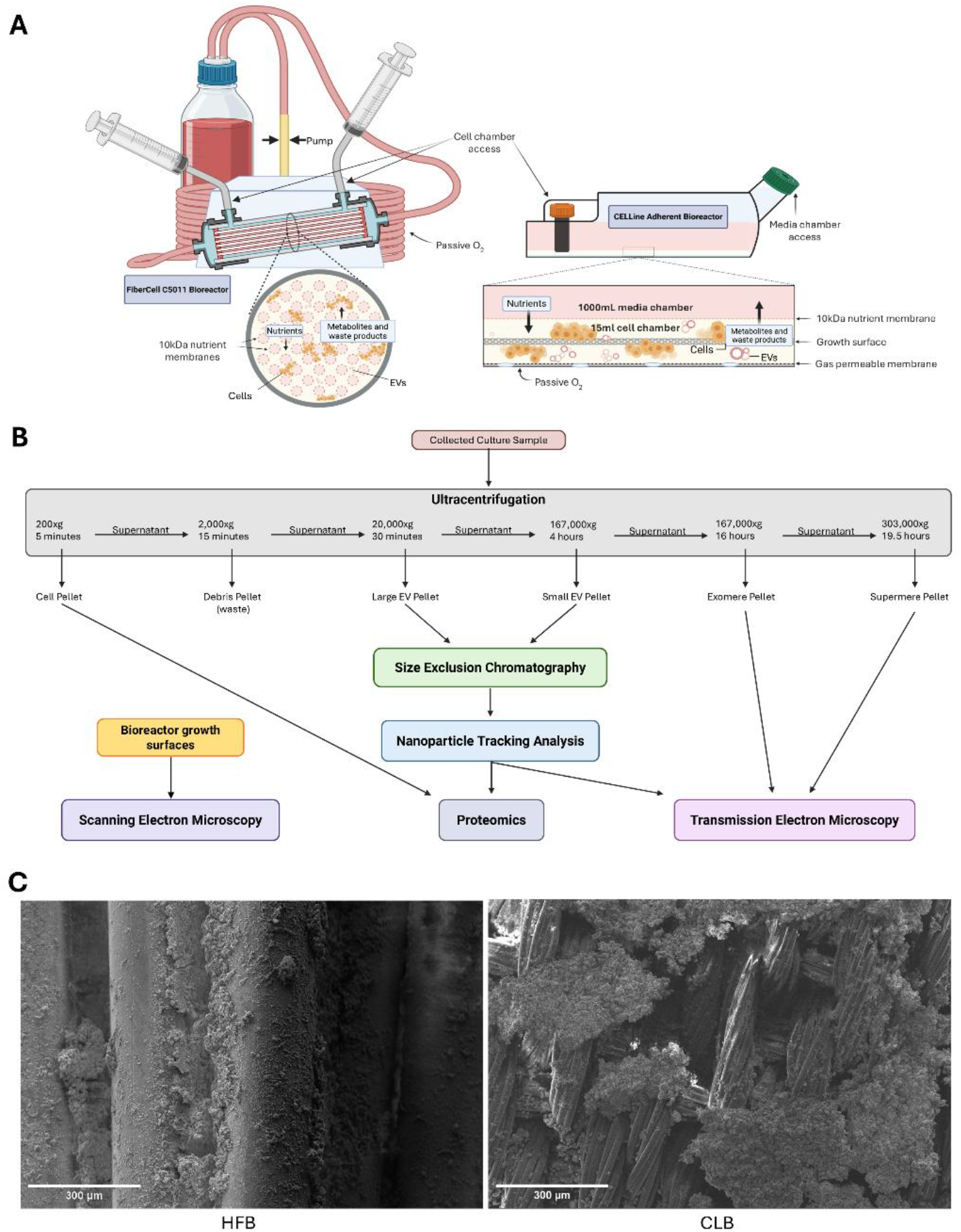
(A) Schematic representations of hollow-fiber and two-chamber bioreactors. Created in BioRender. Tripp, T. (2026) https://BioRender.com/hfncof7 (B) Experimental workflow summarizing EV isolation, purification, and characterization. (C) Representative SEM images of the HFB and CLB growth surfaces. The distinct surface architectures highlight the different culture environments in each system. SEM images at additional magnifications are provided in Supplementary Fig. S1.

## Materials and Methods

### Bioreactor Inoculation, Adaptation, and Maintenance

M1: DMEM with 10% fetal bovine serum (FBS) and 1% antibiotic-antimycotic (anti-anti) M2: DMEM with 5% FBS, 5% CDM-HD (FiberCell Systems), and 1% anti-anti M3: DMEM with 10% CDM-HD and 1% anti-anti.

EVs were isolated from HEK293T cells cultured in a CELLine 1000 AD bioreactor (CLB, WHEATON), a C5011 hollow-fiber bioreactor (HFB, FiberCell Systems), and 175 cm^2^ vented CFs (Thermo Fisher Scientific).

The CLB cell chamber was inoculated with 15 mL of cell suspension (M1, 6.7×10^6^ cells/mL) and its media chamber was filled with 500 mL M1. The HFB cell chamber was inoculated with 20 mL of cell suspension (M1, 10.0×10^6^ cells/mL), its media reservoir was filled with 1000 mL M1, and media circulation was maintained at 150±10 mL/min. The cell chambers in both bioreactors were refreshed every 3–4 days (sample collection), and the media chambers were refreshed every 7 days. To support initial cell attachment and ease cell-transition to serum-free media, M2 was substituted for M1 7 days post-inoculation. M3 was substituted for M2 seven days thereafter. In conjunction with the transition to M3, the CLB media chamber was filled to 800 mL, while the HFB media reservoir was maintained at 1000 mL. Bioreactor cultures were then maintained for 14 weeks before fixation and imaging.

### Conventional Cell Culture as a Comparison

Conventional culture was performed in triplicate, with each replicate consisting of 6 CF, each seeded with 25 mL of cell suspension (M1, 2.1×10^6^ cells/mL). M2 was substituted for M1 24 hours after seeding, and M3 was substituted for M2 24 hours thereafter. The culture medium was then collected 48 hours later for downstream processing.

### Scanning Electron Microscopy

Bioreactors were fixed in a 2% paraformaldehyde + 2.5% glutaraldehyde PBS solution for 20 hours at 4°C. The reactors were then deconstructed using a Dremel rotary tool and approximately 5 mm long growth-surface samples were excised and washed twice with PBS. Samples were post-fixed with 1% osmium tetroxide + 1.5% potassium ferrocyanide in PBS for 40 minutes on ice. The samples were subsequently again washed twice with PBS, and dehydrated by way of 30, 50, 70, 90, 100, and 100% ethanol baths (10 minutes each). Samples were next placed in a 50% ethanol + 50% hexamethyldisilane (HMDS) solution bath, followed by a 100% HMDS bath (10 minutes each), and were then left in a fume hood for complete drying. Fully fixed and dried samples were finally sputter coated with 20 nm gold (Cressington 208HR Sputter Coater system) and then imaged using a Quanta 650 Scanning Electron Microscope (FEI) at 10 kV.

### EV Isolation and Purification

Each extraction was first centrifuged at 200 ×*g* for 5 minutes to obtain a cell pellet for downstream immunoblotting. The supernatant was next centrifuged at 2000 ×*g* for 15 minutes to remove large debris. The clarified supernatant was collected again and processed by differential ultracentrifugation to separate EV populations as well as non-vesicular extracellular nanoparticles: 20000 ×*g* for 30 minutes pelleted large EVs (LEVs), 167000 ×*g* for 4 hours pelleted small EVs (SEVs), 167000 ×*g* for 16 hours pelleted exomeres, and 303000×*g* for 19.5 hours pelleted supermeres (Figure 1C).

Following ultracentrifugation, all crude EV pellets were resuspended in 1 mL Dulbecco’s PBS and were subsequently purified using Sepharose CL-6B resin columns (Apex 6B, Everest Biolabs) for automated size-exclusion chromatography (SEC) (Ascent, Everest Biolabs). EV-enriched fractions were pooled into a single, 1.5 mL purified collection in LoBind tubes (Eppendorf).

### Nanoparticle Tracking Analysis

The size distribution and concentration of SEC-purified EV samples were measured using a PMX-120 ZetaView Monolaser Nanoparticle Tracking Analyzer (NTA) equipped with a 520 nm laser and a CMOS camera. Prior to analysis, EV samples were diluted in PBS to achieve optimal measurement-concentrations (1×10^7^–1×10^8^ particles/mL). For each measurement, videos were recorded at 11 distinct positions along the microfluidic channel within the instrument. The camera sensitivity was set to 81.0 and 83.0, and the shutter speed was 110 and 100, for LEVs and SEVs, respectively, and data were analyzed using ZetaView software version 8.06.01.

For each sample type, particle counts obtained from NTA were converted to total particles per collection. Weekly total yields for each bioreactor were treated as independent observations (n = 14), and CF observations were derived from replicate measurements as described above. Normality of the particle-count distributions within each condition was evaluated using the Shapiro–Wilk test. Because group sizes were modest and variances were not assumed to be equal, overall differences among CLB, HFB, and CF were assessed using the non-parametric Kruskal–Wallis test, followed by post-hoc pairwise comparisons using the Games–Howell test. In addition to hypothesis testing, descriptive statistics were computed for both total particles per collection and particles per milliliter of collected medium for each condition. All statistical analyses and visualizations were performed in Python using SciPy and associated open-source packages, with a significance threshold of α = 0.05.

### Immunoblotting

EV samples from CF, CLB, and HFB were separately pooled and concentrated using ultrafiltration (10 kDa MWCO Sartorius Vivaspin 500) at 10000 ×*g* for 15-minute increments until 100 μL retentate remained. Cells from all three conditions were pooled together and concentrated following the same steps. Concentrated EV and cell samples were then lysed using RIPA lysis buffer, and protein quantities were measured via high-sensitivity BCA (Pierce™ BCA Protein Assay Kit, Thermo Fisher Scientific). Normalized protein amounts (3 μg per lane) were separated using gel electrophoresis at 180 V for 45±10 minutes at 4°C by way of SDS-PAGE (Novex 4–20% Tris-Glycine Plus WedgeWell Gel, Thermo Fisher Scientific), and then transferred to nitrocellulose membranes (Thermo Fisher Scientific) using 10 V over 60 minutes at 4°C.

Following the transfer, membranes were submerged and gently rocked in 20 mL of 0.2 μm filtered 5% BSA in TBS-T for 1 hour at room temperature (RT) to prevent nonspecific binding. Membranes were immunoblotted with GAPDH (Abcam ab9485, 1:1000, reducing), GRP94 (Abcam ab3674, 1:1000, reducing), and CD81 (Abcam ab79559, 1:1000, non-reducing) antibodies for a minimum of 12 hours at 4 °C, followed by either biotinylated anti-rabbit (Jackson Immuno Research, 111065046, 1:10000) or anti-mouse (Jackson Immuno Research, 115065071, 1:10000) secondary IgG antibodies for 1 hour at RT, followed lastly by HRP-conjugated streptavidin (Jackson Immuno Research, 016030084, 1:10000) for 1 hour at RT. Bound antibody was visualized using SuperSignal™ West Pico PLUS Chemiluminescent Substrate (Thermo Fisher Scientific) and imaged using an Azure 400 (Azure Biosystems).

### Label-Free Proteomic Analysis (LC-MS/MS)

Proteins were isolated using SDS-PAGE methodology (38). Briefly, electrophoresis was performed on 20 μg of protein, for each lysed EV sample, at 120 V for 10 minutes to induce short migration and protein separation. Gels were then stained with Coomassie blue, and subsequent visible bands were excised, submerged in ultrapure water, and submitted to the Northwestern University Proteomics Core Facility for mass spectrometry (MS) processing.

Data acquisition was performed using an Orbitrap Exploris 240 Mass Spectrometer (Thermo Fisher Scientific) in data-independent acquisition (DIA) mode over 60 minutes. Full MS spectra were acquired over the range 350–950 m/z at a resolution of 120K. Fragmented-ion spectra were acquired using 75 discrete DIA windows– each spanning 8 m/z– at a resolution of 60K.

Raw DIA data were processed to yield a normalized protein-level intensity matrix, which was analyzed using Python. Intensities were preprocessed per sample by left-censor imputation followed by log2(x+1e−6) transformation, followed by row-wise Z-scaling for visualization. Unsupervised clustering heatmaps were generated from these Z-scores (proteins as rows, samples as columns). Differential variability across experimental groups (CLB-SEV, HFB-SEV, CF-SEV, CLB-LEV, HFB-LEV, CF-LEV) was assessed using one-way, protein-wise ANOVA with Benjamini–Hochberg false-discovery control. Proteins with FDR≤0.05 were retained for downstream analysis. Ordinations (PCA, t-SNE) were fit to the common-protein matrix, with adaptive small-sample safeguards (perplexity cap) and seed-stability checks to flag unreliable ordination. Gene Ontology analysis of the ANOVA-selected list used DAVID; tables were standardized, an adjusted p-value (FDR) was taken, and significance was visualized as −log10(FDR), with terms plotted when significant. Finally, to benchmark EV signal capture, we tested per-sample over-representation against two independent “Top 100 EV protein” references (Vesiclepedia, ExoCarta) using a right-tail hypergeometric model with BH correction, reporting overlap counts, expected overlaps, enrichment, and adjusted p-values.

### Transmission Electron Microscopy

Droplets of EV samples (20 µL) were adsorbed onto carbon Type-B film supported 300 mesh copper grids for 20 minutes (Ted Pella Inc.). The grids were then washed twice with ultrapure water (10 µL, 30 seconds), and negative stained with 10 µL UranyLess for 30 seconds (Electron Microscopy Sciences). Between each step, excess liquid was blotted with filter paper (Whatman). Grids were airdried overnight, before being imaged using a JEM-1400Flash Transmission Electron Microscope (JEOL) at 120 kV.

### PEG-PPS Synthesis and Characterization

PEG-PPS-Bz was synthesized as previously described (39–42). Briefly, for MC, mPEG45-OH (Sigma-Aldrich) was functionalized with mesylate groups followed by thioacetic acid to yield mPEG45-thioacetate. mPEG45-thioacetate was then reacted with sodium methoxide (Sigma-Aldrich) to generate a thiolate anion followed by addition of propylene sulfide (Tokyo Chemical Industry) and end capped with benzyl bromide (Sigma-Aldrich) to yield PEG_45_-PPS_21_-Bz. Identity of all the products were confirmed using ^1^H-NMR (see Supplementary Fig. S4).

### Nanocarrier Formulation and Characterization

Micelles were formed by using thin film hydration. Briefly, 20 mg of PEG-PPS-Bz polymer was weighed into a glass vial, dissolved in organic solvent, and evaporated to form thin films. For MCs loaded with Cy3 (MC-L-Cy3), 2.5 mol% Cyanine3-carboxylic acid (Lumiprobe) was dried with the thin film. After desiccation for several hours, 400 µL of 1x PBS was added to the films and vigorously shaken overnight. MCs were then purified by using 7K MWCO Zeba Desalting Columns (Thermo Fisher Scientific) followed by dialysis in 7K MWCO cassettes (Thermo Fisher Scientific) for 2 days to remove excess Cyanine3-carboxylic acid.

For MCs grafted with AlexaFluor555 (MC-G-AF555), 2.5 mol% N_3_-PEG-1,2-Distearoyl-sn-glycero-3-Phosphoethanolamine (N_3_-PEG-DSPE) was added and continued to be vigorously shaken overnight. Nanocarriers were then purified using 7K MWCO Zeba Desalting Columns (Thermo Fisher Scientific) to remove excess N_3_-PEG-DSPE. N3-PEG-DSPE was then reacted with 1.1 eq AlexaFluor 555-Dibenzocyclooctyne (AF555-DBCO, Lumiprobe). Excess AF555-DBCO was removed using 7K MWCO Zeba Desalting Columns followed by dialysis in 7K MWCO cassettes for 2 days to remove excess AF555-DBCO.

After, formulations were filtered multiple times using 0.2 µm filters. Dynamic light scattering (Malvern Instruments) was used to verify size, monodispersity, and zeta potential.

### EV Engineering

Size distribution and concentration of purified EVs were first assessed as described above to establish a baseline. Two engineering procedures were utilized to yield cargo-loaded and surface-modified EVs, wherein EVs were loaded with Cy3 fluorophores (EnEV-L-Cy3) and were grafted with AlexaFluor555 conjugated PEG-lipids (EnEV-G-AF555) respectively.

For cargo-loading, 200 µL of EV suspension was combined into 0.5 LoBind tubes with 75 µL of 2 mg/mL MC-L-Cy3 suspension and 25 µL PBS (in triplicate). Tubes were sonicated for 30 seconds followed by 2 minutes of rest at 10°C for 12 cycles followed by 1 hour incubation at 37°C, and lastly 15 minutes incubation on ice.

For surface-modification, 200µL of EV suspension was combined into 0.5 LoBind tubes with 75 µL of 2 g/mL MC-G-AF555 suspension and 25 µL PBS (in triplicate). Tubes were incubated at 37°C for 2 hours, followed by 15 minutes incubation on ice (43,44).

## Results

### EV Production Yields Across Conditions

Per collection, it was observed that CLB and HFB statistically outperformed CF in LEV production (p<0.01). For SEV production, HFB was observed to statistically outperform CF (p<0.05), while no statistical difference was observed between CF and CLB (Figure 2A). While large variability in EV yield was noted across timepoints for both bioreactor types, they consistently remained at or above CF yields (Figure 2B). Additionally, size distributions of bioreactor-derived EVs remained stable across timepoints as shown in Figures 2C and 2D. Differences in performance were determined by normalized yield per mL of media, with means of 2.81×10^10^, 6.39×10^10^, and 2.19×10^8^ particles/mL for CLB, HFB, and CF respectively.

**Figure 2.**
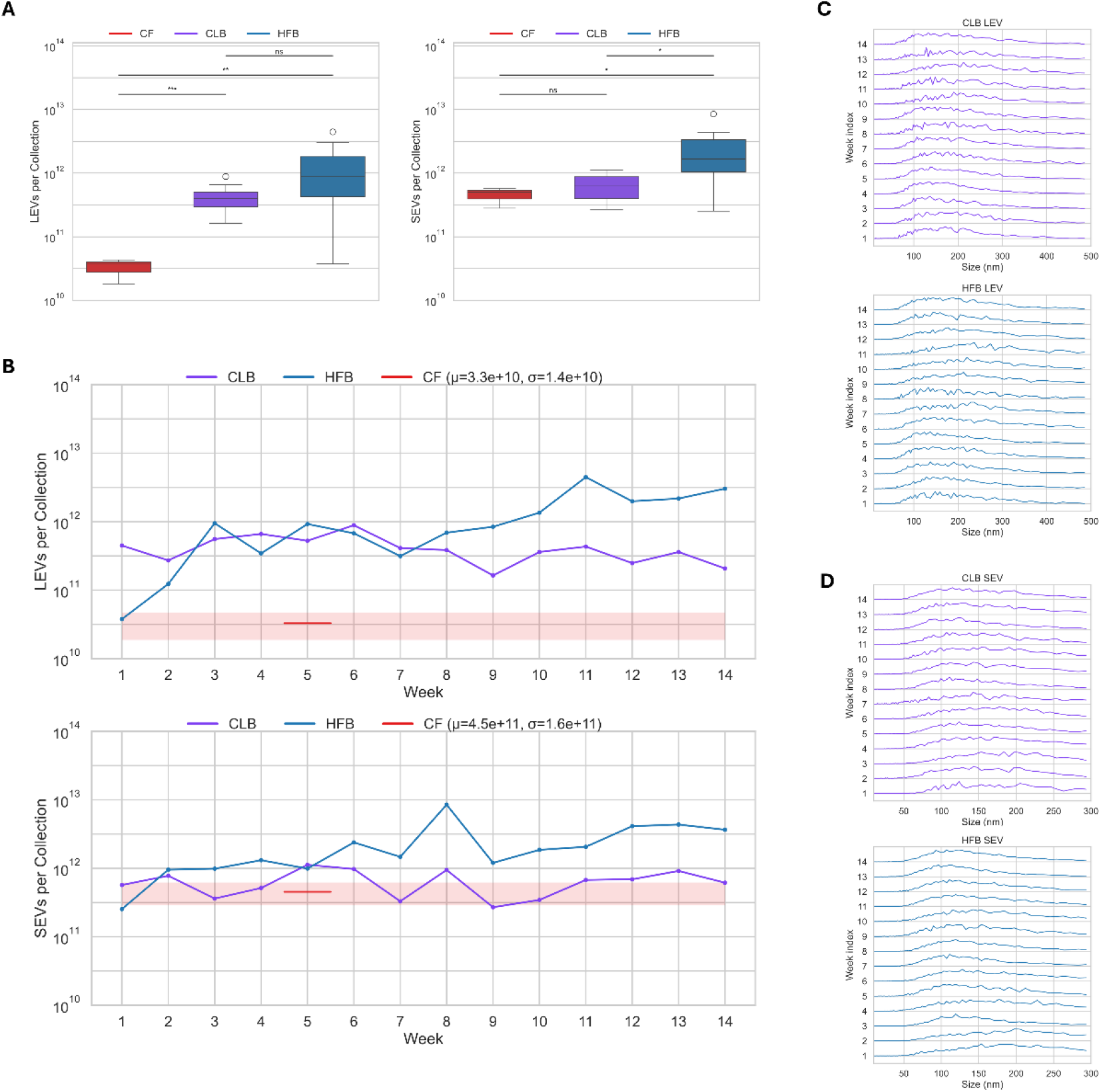
(A) Box plots comparing LEV and SEV production per harvest across CF, CLB, and HFB. (B) Longitudinal production profiles for LEVs and SEVs over the course of the study. (C,D) Size distribution ridge plots of all collected EVs binned by week.

### EV Identification and Proteomic Profiling

Immunoblotting resulted in visible band-appearance in each of the expected regions for both LEV- and SEV-blots. The ∼94 kDa bands corresponding to GRP94 were not detected in EV samples, with faint detection in cell lysate. Visibility of the GRP94 bands was augmented with a uniform increase in image brightness and contrast. With the exception of HFB-SEV, bands corresponding to GAPDH at ∼37 kDa were visible in all samples. Bands at ∼22 kDa corresponding to CD81 were observed in all EV samples and were absent from cell lysate. Full, uncropped immunoblot membrane images are shown in Supplementary Fig. S2.

Protein mass spectrometry analysis revealed strong overlap across CF, CLB, and HFB for both LEVs and SEVs through the first three weeks of collections (Figure 3B). Gene expressions across all samples are shown in Figure 3C. Notably, CLB and HFB samples appear largely distinct from CF samples apart from two early-timepoint samples. Principle component analysis (PCA) similarly resulted in bioreactor samples appearing predominantly distinct from CF samples, and subsequent t-distributed stochastic neighbor embedding (t-SNE) followed a similar trend (Figure 3D). These observed phenotypic differences were further investigated through GO analysis of the most variably expressed proteins, which found statistically variable pathways across samples belonging exclusively under Cellular Component and were related to metabolism (Figure 3E). TEM images shown in Figure 3F capture expected EV morphology. Of note, bioreactors were observed to also produce supermeres and exomeres (Figure 3G).

**Figure 3.**
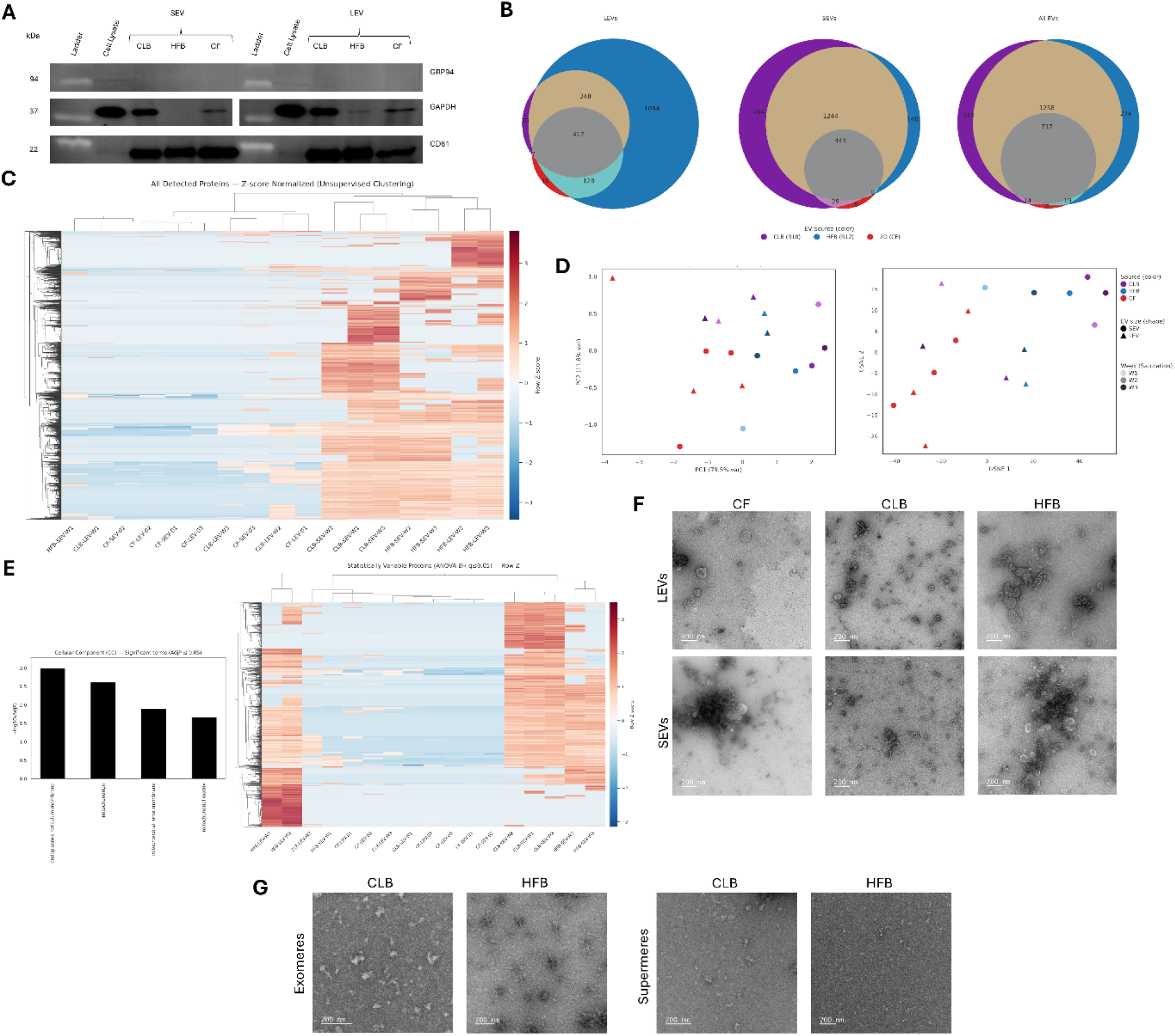
(A) Immunoblot analysis of canonical EV protein markers for LEVs and SEVs collected from all conditions. (B) Venn diagrams illustrating overlapping and unique proteins identified in LEV, SEV, and combined fractions across all conditions. (C) Heatmap of normalized protein intensities for samples across the first three weeks of production. Unsupervised clustering shows grouping by each platform. (D) PCA and t-SNE plots of proteomic profiles show distinction of EV populations by culture systems. (E) Heatmap of the most variable proteins across all samples accompanied by a bar chart of significantly enriched pathways derived from these proteins. (F) Representative TEM images of LEVs and SEVs from each platform, demonstrating characteristic EV morphology. (G) Representative TEM images of exomeres and supermeres isolated from bioreactor platforms. See Supplementary Fig. S3 for additional TEM images at lower magnification.

### EV Engineering

Schematic representations of the two engineering approaches are shown in Figure 4A-B. MC characterization revealed particle size distributions far lower than the approximate 50 nm NTA detection limit as shown in Figure 4B (45,46). Experimental processing was observed to have negligible impact on EV size distribution, with small increases in counted particles correlating to micelle detection (Figure 4C-D). Imaging of the experimental groups demonstrated a lack of MC aggregation and potential hybridization between MCs and EVs. NTA readings for MCs and EVs prior to engineering procedures reported zero fluorescent particles per mL for both experimental conditions. Following experimentation, EV and MC controls remained non-fluorescent, while fluorescence was observed in both engineered EV samples (Figure 4E-F).

**Figure 4.**
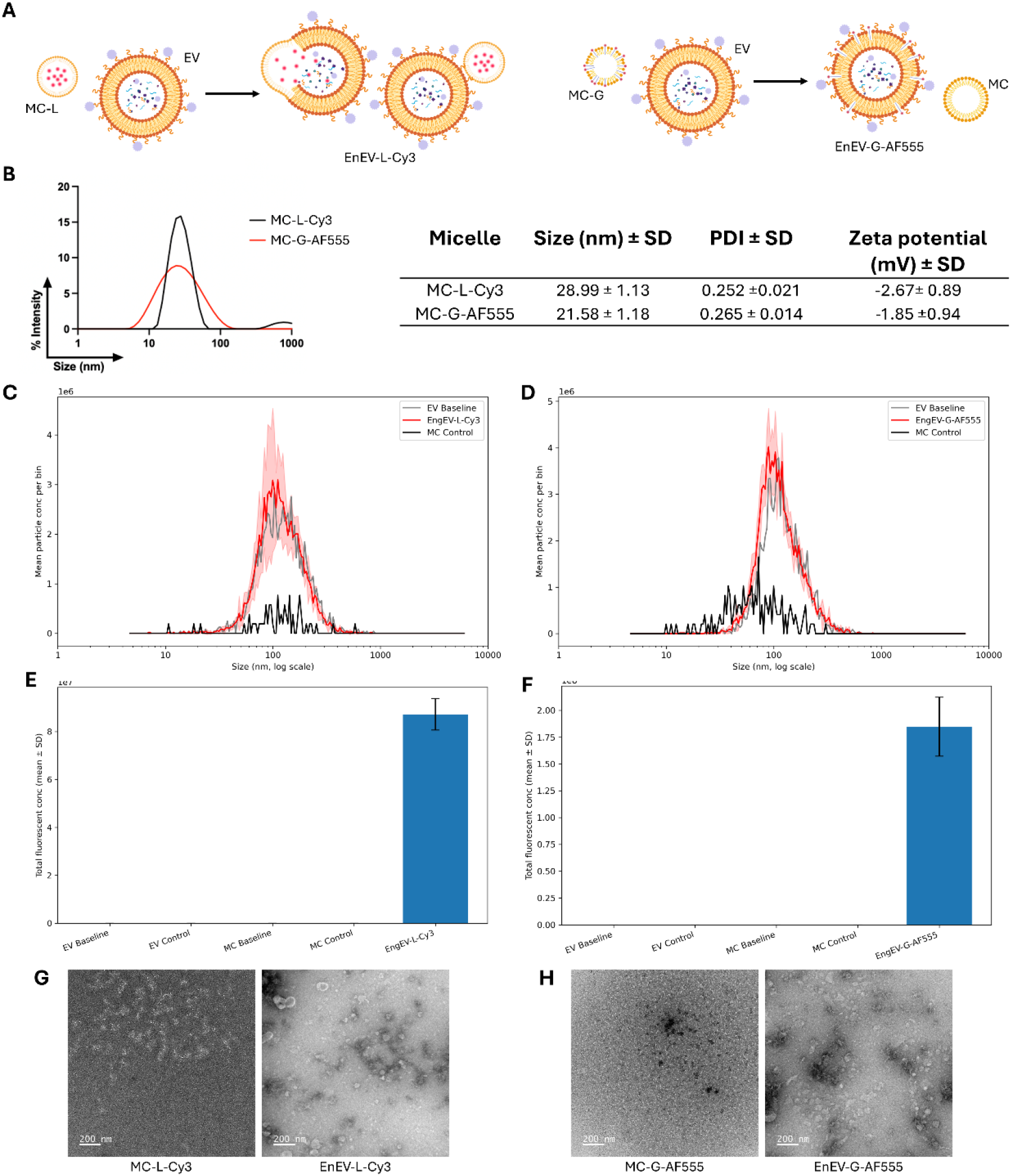
(A) Schematic illustrations of cargo-loading and surface-modification engineering showing expected hybrid formations. Micelles loaded with Cy3 (shown as red particles) fuse with EVs, and micelles with membrane-inserted AlexaFluor555 PEG-lipids donate fluorophores to EVs via post-insertion. Created in BioRender. Patel, A. (2026) https://BioRender.com/km2jl1r (B) Size distribution of Cy3-loaded micelles and AlexaFluor555-grafted micelles with accompanying summary table of micelle characteristics. (C,D) Size distribution curves for engineered EVs and corresponding EV and micelle controls post-engineering. Plots show particle-number increases in experimental conditions relative to baseline. (E,F) Quantification of fluorescence signals following engineering demonstrate incorporation of micelle-associated fluorescence. (G,H) Representative TEM images of micelles and engineered EVs for cargo-loading and surface-modification engineering.

## Discussion

HFBs produced substantially higher EV yields relative to CFs, reinforcing prior reports that high cell density bioreactor systems can increase EV output by orders of magnitude (25,33). These findings support the view that HFBs are particularly well suited for scalable EV manufacturing. In concert with growing evidence that engineered EVs can mediate tissue repair and foster diverse therapeutic effects (18,47), our data position HFBs as a practical backbone for future cGMP-oriented EV production.

The bioreactor systems also demonstrated several operational advantages that directly impact translational feasibility. Compared to equivalent cell expansion in flasks, bioreactors required markedly less media per EV produced, reducing both reagent consumption and waste. This is consistent with previous work demonstrating reduced resource burden associated with high-density culture conditions (25,33). Furthermore, the semi-closed design of these systems reduces contamination risk and aligns with manufacturing principles emphasized in EV standardization and translational guidelines (48). In combination with the observed data, these principles suggest that bioreactors are intrinsically more amenable to clinical translation than traditional culture flasks.

EV identity was validated through immunoblotting, demonstrating robust EV production in bioreactors. However, band intensities for select markers varied across replicates, and we lacked a guaranteed loading control for certain populations (HFB SEVs), mirroring a broader challenge in the field. The absence of a universally accepted EV housekeeping protein and the sensitivity of EV marker detection to sample preparation, antibody choice, and platform-specific variables have been widely noted (38,48,49). Our data therefore underscores the importance of using multiple orthogonal characterization metrics (size distribution, morphology, and marker panels), along with persistent efforts toward assay and reporting standardization.

Despite these analytical challenges, the phenotype of bioreactor-derived EVs remained conducive to translational applications. The observed differences in proteomics profiles likely reflect the 3D culture environment rather than alterations of EV identity associated with cellular disfunction. Given that bioreactors expose cells to nutrient-, oxygen-, and waste-gradients that differ from culture flasks, and such factors are known to modulate cargo loading (25,33), the phenotypic shifts we observed should be interpreted as bioreactor-specific tuning of EV profiles– potentially exploitable for application-specific optimization. Additionally, we have demonstrated that bioreactors can be used to produce exomeres and supermeres at scale, which could offer another avenue for engineered nanotherapeutics (50–53).

A central motivation of this work was to determine whether the enhanced yields achieved with bioreactors translate into more practical EV engineering. Here, we demonstrate that all engineering experimentation combined consumed less than 8% v/v of all purified EVs produced by a given bioreactor over the 14-week lifetime of the study. This scale would be difficult to achieve using culture flasks alone. This is particularly important because many commonly used engineering methods are intrinsically low-efficiency (15). Our data show that, when coupled to high-yield bioreactor production, even these inefficient approaches become operationally feasible, enabling sufficient engineered EV material for characterization and downstream testing.

Crucially, bioreactor-derived EVs showed no detectable inherent limitations in their engineerability. Previous studies have established that CF-derived EVs can be engineered (54–58), and our data indicate that bioreactor-derived EVs likewise maintain compatibility with incubation- or sonication-based engineering protocols. Instead, the main bottleneck in engineered EV yield appears to be the absolute quantity of EVs available for modification, which is directly alleviated by bioreactors.

It is worth highlighting that the employed engineering methodology exploited the limitations of the NTA instrument to validate successful EV engineering. Because the 20-30 nm fluorescent MCs were outside the detection range of the instrument, no post-engineering clean-up step was required before assessing hybridization. This meant that any fluorescence observed was the result of either EV-MC hybridization or MC aggregation. Given the lack of fluorescence present in the MC controls, we can confidently conclude that both EV engineering modalities were successful. It should be emphasized that, for the cargo-loading procedure (Figure 4A), no statement is being made on the respective prevalences of the potentially different forms or structures of the final hybrid nanoparticles. This detection technique could be further leveraged for high-throughput screening of hybridization success across differently formulated MCs to efficiently optimize EV engineering yield. For downstream clinical use, the large difference in size between EVs and MCs can again be leveraged using SEC for quick and simple post-engineering clean-up of excess MCs. Moreover, we observed, to the best of our knowledge, the first experimental evidence of EV-MC hybridization. This provides a technical foundation for future development of hybrid EV systems.

Collectively, these findings highlight that bioreactors not only overcome the throughput limitations of flask-based production but also generate phenotypically suitable, readily engineerable EV populations. Future studies should expand this framework to additional cell types, longer production windows, and more diverse engineering modalities.

## Conclusion

In this study, we systematically compared EV production from conventional culture flasks and bioreactor systems. We demonstrated that hollow-fiber systems drastically increase EV yield while simultaneously reducing operational burden and highlight the strong potential of such platforms for cGMP-oriented manufacturing. These findings are consistent with prior work showing that high-density bioreactor cultures can achieve substantially higher EV productivity than monolayer systems.

We further show that EVs harvested from bioreactors retain canonical phenotypic features and engineering efficacy, even when using low-efficiency methods such as passive incubation and sonication. Because engineering experiments consumed only a small fraction of the total EV yield, our results directly address limited availability of starting material. Bioreactor-derived EVs therefore offer a powerful means to convert existing engineering techniques into scalable workflows. Together, these data support a model in which bioreactor platforms, particularly hollow-fiber systems, serve as a translational bridge between laboratory-scale EV research and clinically relevant production.

## Acknowledgements

This work made use of the EPIC facility (RRID: SCR_026361) and the BioCryo facility (RRID:SCR_021288) of Northwestern University’s NUANCE Center, which has received support from the SHyNE Resource (NSF ECCS-2025633), the IIN, and Northwestern’s MRSEC program (NSF DMR-2308691). Proteomics services were performed by the Northwestern Proteomics Core Facility (RRID:SCR_017945), generously supported by NCI CCSG P30 CA060553 awarded to the Robert H Lurie Comprehensive Cancer Center, instrumentation award (S10OD025194) from NIH Office of the Director, and the National Resource for Translational and Developmental Proteomics supported by P41 GM108569. This work was supported by the National Institutes of Health grants: C.L.H. supported by R00EB033857 and E.C. supported by T32T32GM153505 (Northwestern University’s Biotechnology Training Program).

## Author Contributions

T.T.: Conceptualization, Methodology, Formal Analysis, Investigation, Writing – Original Draft, Visualization. K.V.: Investigation, Writing – Original Draft, Writing – Review & Editing. M.T.: Investigation, Writing – Original Draft, Writing – Review & Editing. E.C.: Methodology, Formal Analysis, Writing - Original Draft. L.V.: Resources, Writing – Review & Editing. C.H.: Conceptualization, Resources, Writing – Review & Editing, Supervision, Funding acquisition.

