## Supplementary for "Bioreactor-Enabled Extracellular Vesicle Production for Downstream Functional Engineering"

**Table of Contents:**

**Supplementary Figure 1.** Additional growth surface imaging using SEM….………………………………………………….……………**S1**

**Supplementary Figure 2.** Immunoblots….……………………………………………………………..………………………………………………….**S2**

**Supplementary Figure 3.** Additional EV imaging using TEM……………………………………………………………….……………..……….**S3**

**Supplementary Figure 2.** ^1^H-NMR of PEG_45_-PPS_21_-Bz….…………………………………………..………………………………………………….**S4**

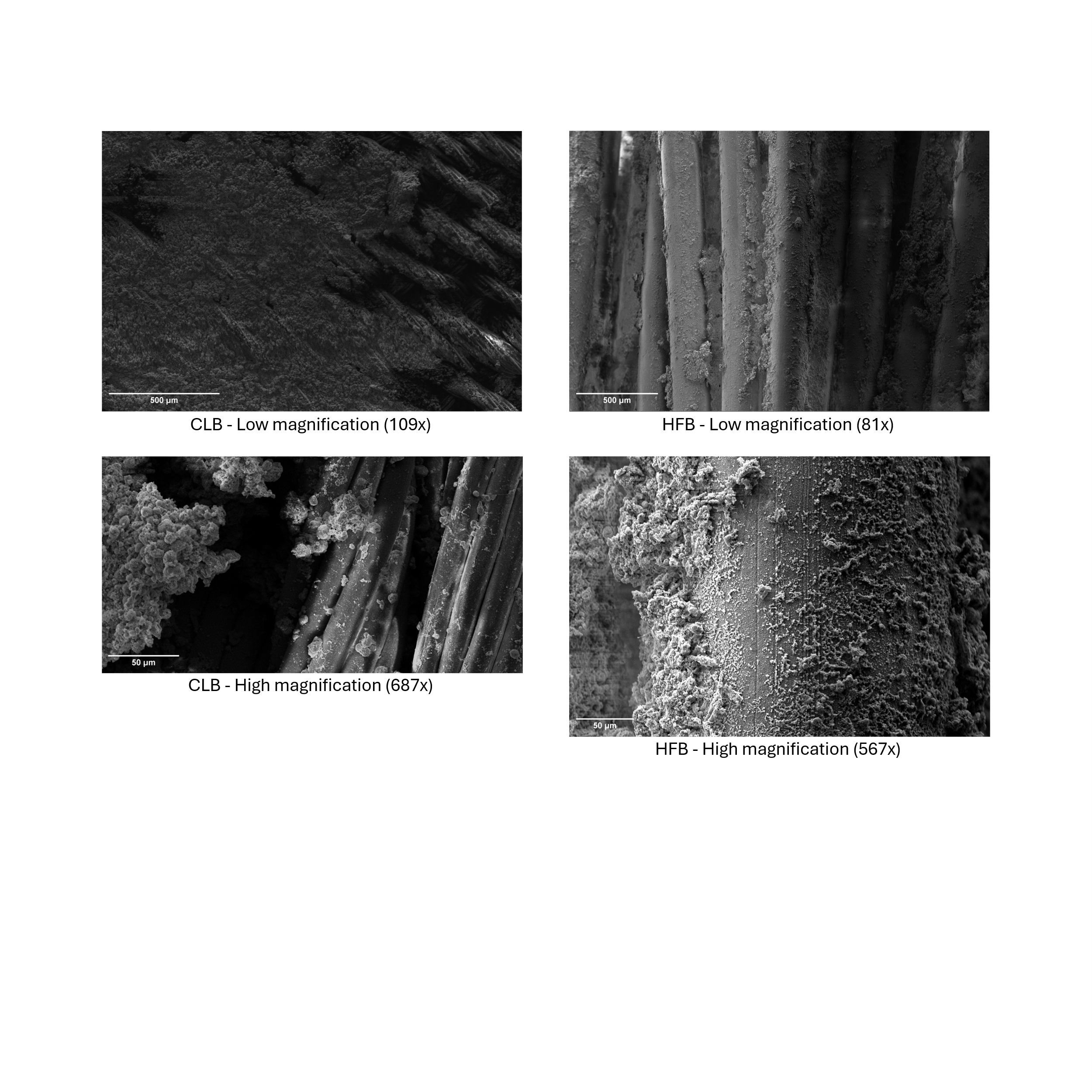

**Supplementary Figure 1.** SEM images of CLB and HFB growth surfaces at low and high magnifications.

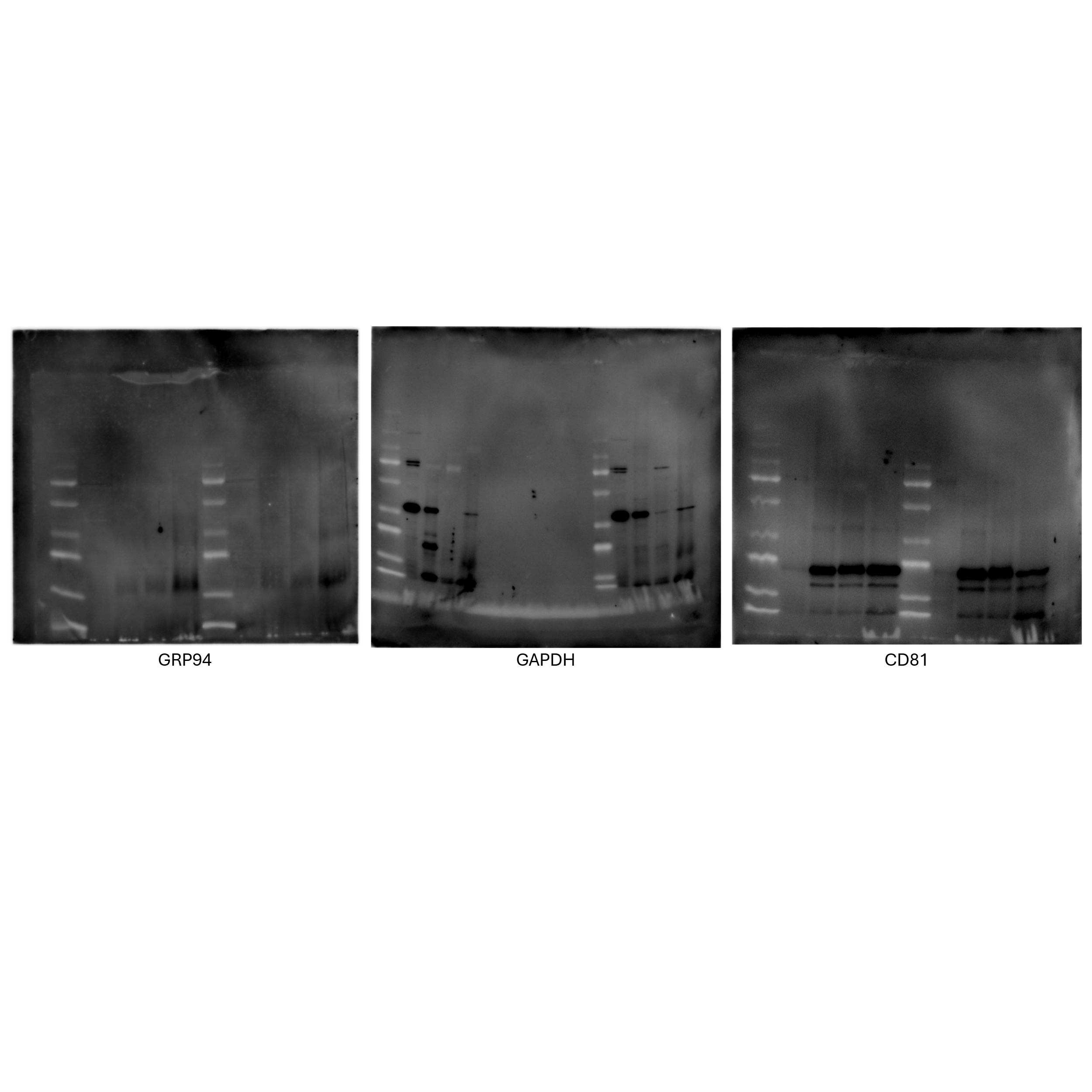
**Supplementary Figure 2.** Full immunoblot membrane images used for the cropped blots presented in the main text.

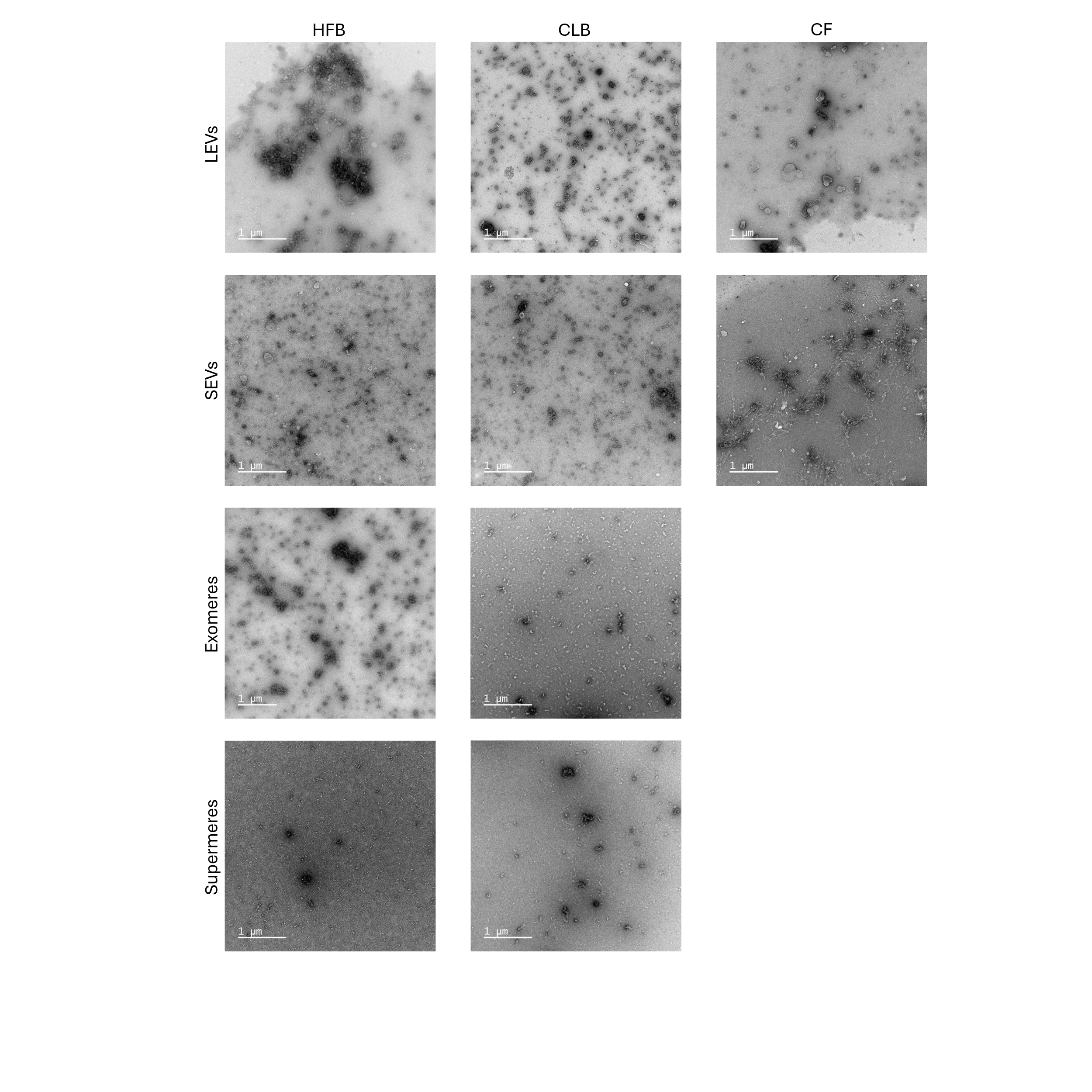

**Supplementary Figure 3.** TEM images of LEVs and SEVs across all growth conditions accompanied by TEM images of bioreactor-derived exomeres and supermeres.

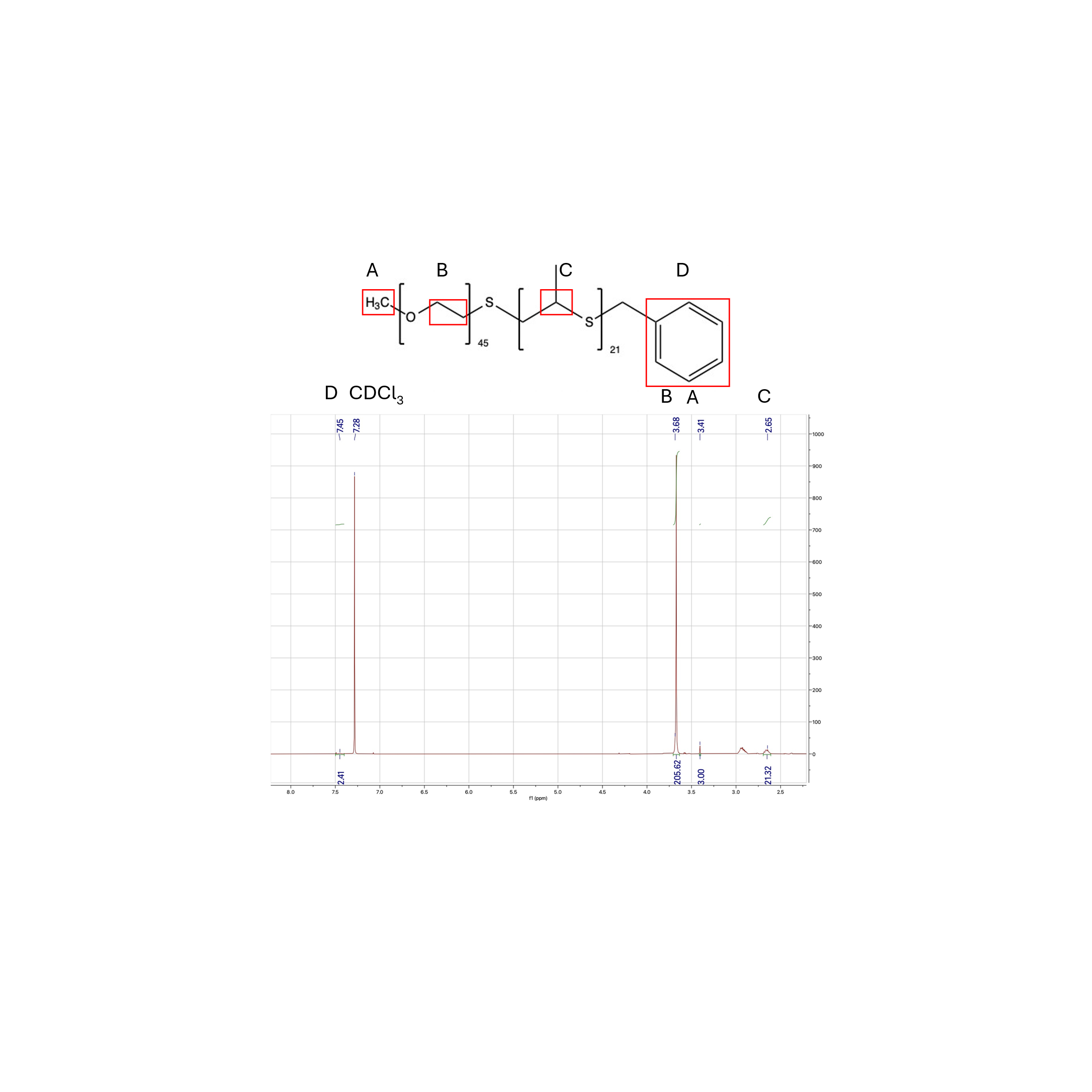

**Supplementary Figure 4.** ^1^H-NMR of PEG_45_-PPS_21_-Bz
